# Differential Activity-Dependent Increase in Synaptic Inhibition and Parvalbumin Interneuron Recruitment in Dentate Granule Cells and Semilunar Granule Cells

**DOI:** 10.1101/2021.05.18.444756

**Authors:** Milad Afrasiabi, Akshay Gupta, Huaying Xu, Bogumila Swietek, Vijayalakshmi Santhakumar

## Abstract

Strong inhibitory synaptic gating of dentate gyrus granule cells (GCs), attributed largely to fast-spiking parvalbumin interneurons (PV-INs), is essential to maintain sparse network activity needed for dentate dependent behaviors. However, the contribution of PV-INs to basal and input driven sustained synaptic inhibition in GCs and semilunar granule cells (SGCs), a sparse morphologically distinct dentate projection neuron subtype are currently unknown. In studies conducted in hippocampal slices from mice, we find that although basal inhibitory postsynaptic currents (IPSCs) are more frequent in SGCs and optical activation of PV-INs elicited IPSCs in both GCs and SGCs, optical suppression of PV-INs failed to reduce IPSC frequency in either cell type. Amplitude and kinetics of IPSCs evoked by perforant path activation were not different between GCs and SGCs. However, the robust increase in sustained polysynaptic IPSCs elicited by paired afferent stimulation was lower in SGCs than in simultaneously recorded GCs. Optical suppression of PV-IN selectively reduced sustained IPSCs in SGCs but not in GCs. These results demonstrate that PV-INs, while contributing minimally to basal synaptic inhibition in both GCs and SGCs in slices, mediate sustained feedback inhibition selectively in SGCs. The temporally selective blunting of activity-driven sustained inhibitory gating of SGCs could support their preferential and persistent recruitment during behavioral tasks.

**Significance Statement:** Our study identifies that feedback inhibitory regulation of dentate semilunar granule cells, a sparse and functionally distinct class of projection neurons, differs from that of the classical projection neurons, granule cells. Notably, we demonstrate relatively lower activity dependent increase in sustained feedback inhibitory synaptic inputs to semilunar granule cells when compared to granule cells which would facilitate their persistent activity and preferential recruitment as part of memory ensembles. Since dentate granule cell activity levels during memory processing are heavily shaped by basal and feedback inhibition, the fundamental differences in basal and evoked sustained inhibition between semilunar granule cells and granule cells characterized here provide a framework to reorganize current understanding of the dentate circuit processing.

## Introduction

The hippocampal dentate gyrus is known for its uniquely sparse activity, essential for its function in pattern separation, a process of disambiguating similar inputs into distinct patterns of neural activity and memory representations (Bekinschtein et al., 2013; GoodSmith et al., 2017). The characteristically sparse dentate activity is maintained by the relatively hyperpolarized resting membrane potential and low input resistance of the projection neurons and their powerful inhibitory regulation (Heinemann et al., 1992; Lothman et al., 1992). Dentate inhibition includes sustained tonic/extrasynaptic GABA currents as well as distinct phases of synaptic inhibition, which shape basal and afferent-evoked activity (Stell et al., 2003; Ewell and Jones, 2010). Apart from the classical basal spontaneous synaptic inhibition and afferent evoked feedforward and feedback inhibition, dentate granule cells (GCs) have been proposed to receive “sustained” increase in feedback synaptic inhibition following paired stimulation of perforant path (PP) inputs (Larimer and Strowbridge, 2010). This sustained feedback inhibition has been suggested as a mechanism for surround inhibition needed to facilitate pattern separation (Larimer and Strowbridge, 2010; Walker et al., 2010). However, the reliability, cell-specificity and temporal structure of this sustained feedback inhibition is not known and the identity of inhibitory neurons recruited during this prolonged feedback inhibition has not been examined.

Although dentate GCs have, for long, been considered a structurally and functionally uniform class of projection neurons, detailed morphometric and physiological analysis in rats have revealed a sparse, structurally distinct population of neurons with axonal projections to CA3 described as semilunar granule cells (SGCs) by Ramón y Cajal (1995) (Williams et al., 2007; Gupta et al., 2012; Save et al., 2018). Functionally, SGCs have sustained firing in response to paired PP stimulation, which distinguish them from GCs (Larimer and Strowbridge, 2010). While SGCs, characterized by wider dendritic span, have been reported in mice (Save et al., 2018; Erwin et al., 2020), detailed morphometric analysis and functional characterization of SGCs in mice is lacking. Of particular relevance to dentate inhibitory regulation, whether SGCs receive similar early (feedforward or disynaptic feedback) as well as sustained polysynaptic feedback inhibition as GCs or may escape this inhibitory gate is currently unknown. Studies in adolescent rats show that SGCs have higher tonic GABA currents and receive more frequent synaptic inhibitory inputs than GCs (Gupta et al., 2012; Gupta et al., 2020). Uniquely, SGCs show reduction in both tonic GABA currents and frequency of synaptic inhibitory inputs after brain injury rather than the increase observed in GCs (Gupta et al., 2012). While these findings suggest different interneuron subtypes may contribute to synaptic inhibition in GCs and SGCs, which class of dentate interneurons underlie this differential control, is not known. Parvalbumin-interneurons (PV-INs) are a major class of GABAergic neurons which are classified by expression of the calcium binding protein parvalbumin and their fast, non-adapting firing pattern. PV-INs include the perisomatically projecting basket cells which contribute to feedforward and feedback inhibition of GCs (Kraushaar and Jonas, 2000; Hefft and Jonas, 2005; Ewell and Jones, 2010) and axo-axonic cells which target granule cell axon-initial segments (Sik et al., 1997; Howard et al., 2005). A single basket cell can project to as many as 10,000 GCs (Freund and Buzsaki, 1996; Santhakumar, 2008) and is an ideal candidate to mediate surround inhibition. However, whether PV-INs innervate SGCs with somata located in the inner molecular layer, outside the dense axon collaterals of parvalbumin positive basket cells has not been tested.

Recent studies have indicated that SGCs may be disproportionately recruited to memory engrams (Erwin et al., 2020) making it important to understand whether inhibitory regulation of SGCs differs from that of GCs. This study was conducted to systematically examine the cell-type specific differences in distinct phases of synaptic inhibition; basal spontaneous synaptic inhibition, PP evoked, early feedforward, and feedback inhibition as well as sustained feedback inhibition, between the two classes of dentate projection neurons and to determine the contribution of PV-INs to specific phases of inhibition in GCs and SGCs.

## Materials and Methods

### Animals

All experiments were performed in accordance with IACUC protocols approved by Rutgers-NJMS, Newark, NJ, and the University of California at Riverside, CA and in keeping with the ARRIVE guidelines. The study included male and female wild-type C57BL/6J (WT) or appropriately validated optogenetic mice (4-8 weeks old, Jackson Laboratory, Bar Harbor, ME). Parvalbumin (PV)-Cre mice (B6;129P2-Pvalbtm1(cre)Arbr/J; JAX #8069) were crossed with either of the two floxed lines (Chr2-YFP: B6;129S-Gt(ROSA)26Sortm32(CAG-COP4*H134R/EYFP)Hze/J; JAX#12569 or NpHR3-YFP: Rosa-CAG-LSL-eNpHR3.0-EYFP-WPRE; JAX #14539) to generate experimental PV-Chr2 and PV-NpHR3 mice. Immunofluorescence labeling for PV and YFP in sections from PV-YFP mice identified that over 80% of PV labeled cells expressed YFP, and all YFP positive cells expressed PV validating the specificity of the PV-Cre transgenic lines. Mice were housed with littermates (4 mice per cage) in a 12-h light-dark cycle. Food and water were provided ad libitum.

### Slice Physiology

Animals were anesthetized under 3% isoflurane and decapitated. Whole brains were extracted and horizontal brain slices (350 µm) were prepared using Leica VT1000 or VT1200S Vibratomes (Wetzlar, Germany) in ice-cold high Sucrose-aCSF containing (in mM) 85 NaCl, 75 sucrose, 24 NaHCO_3_, 25 glucose, 4 MgCl_2_, 2.5 KCl, 1.25 NaH_2_PO_4_, and 0.5 CaCl_2_. Slices were bisected and incubated at 32 ± 1°C for a minimum of 30 min in a submerged holding chamber containing recording aCSF and subsequently held at room temperature (RT, 22-23°C). The recording aCSF contained (in mM) 126 NaCl, 2.5 KCl, 2 CaCl_2_, 2 MgCl_2_, 1.25 NaH_2_PO_4_, 26 NaHCO_3_ and 10 D-glucose. All solutions were saturated with 95% O_2_ and 5% CO_2_ and maintained at a pH of 7.4 for 2-6 h (Gupta et al., 2012; Yu et al., 2016).

Slices (350 µm) were transferred to a submerged recording chamber and perfused with oxygenated aCSF at 33 ± 1°C. Whole-cell voltage- and current-clamp recordings from GCs, SGCs and presumed interneurons at the border of the hilus and GCL were performed under IR-DIC visualization with a Nikon Eclipse FN-1 (Nikon Corporation, Shinagawa, Tokyo, Japan) or Olympus BX50 (Olympus Corporation, Shinjuku, Tokyo, Japan) microscope, using 40X water-immersion objectives. Recordings were obtained using Axon Instruments Axopatch 200B or MultiClamp 700B amplifiers (Molecular Devices, Sunnyvale, CA). Data were obtained using glass microelectrodes (3–5 MΩ) pulled using a Narishige PC-10 puller (Narishige Japan, Tokyo Japan) or Sutter P-1000 Glass puller (Sutter Instruments, Novato, CA) and were low-pass filtered at 3 kHz, digitized using DigiData 1440A and acquired using pClamp10 at 10-kHz sampling frequency with gains of 0.1 and 1. Recordings were performed using high chloride internal solution containing (in mM) 125 KCl, 10 K-gluconate, 10 HEPES, 2 MgCl_2_, 0.2 EGTA, 2 Na-ATP, 0.5 Na-GTP and 10 PO Creatine titrated to a pH 7.25 with KOH (pH 7.25; 270–290 mOsm) or cesium methane sulfonate (CsMeSO_4_) internal solution containing (in mM) 140 cesium methane sulfonate, 10 HEPES, 5 NaCl, 0.2 EGTA, 2 Mg-ATP, 0.2 Na-GTP (pH 7.25; 270–290 mOsm). Biocytin (0.3%) was included in the internal solution for post-hoc cell identification (Gupta et al., 2012; Yu et al., 2015). In experiments with KCl based internal, synaptic GABA currents were recorded in perfusing aCSF containing the glutamate receptor antagonist kynurenic acid (3 mM, Tocris, Ellisville, MO). Recorded neurons were initially held at −70 mV for interneurons and −60 mV for GCs and SGCs and the response to 500 milli-second positive and negative current injections were examined to determine active and passive characteristics. Voltage Clamp recordings of IPSCs (spontaneous and evoked) were obtained from a holding potential of −70 mV for interneurons and −60 mV for GCs and SGCs (Yu et al., 2015; Gupta et al., 2020). CsMeSO_4_ based internal was used to record the IPSCs as outward currents from a holding potential of at 0 mV in the absence of glutamate receptor antagonists. In some experiments tetrodotoxin (TTX, 1 µM) or gabazine (10 µM) were used to isolate action potential independent miniature IPSCs or block IPSCs, respectively. Post-hoc biocytin immunostaining and morphological analysis was used to definitively identify interneurons, GCs and SGCs included in this study. Morphologies were reconstructed using Neurolucida360 (Microbrightfield) for further analysis (See Morphometry below).

Evoked IPSCs were elicited by single or paired stimulation of the PP using bipolar concentric stimulating electrodes placed at the junction of the dorsal blade and the crest, just outside the fissure (Korgaonkar et al., 2020). Constant current stimuli (0.5 μA-5 mA) were applied using a high-voltage stimulus isolator (A365R, World Precision Instruments, Sarasota, FL). The evoked mono and polysynaptic IPSCs were recorded simultaneously in GCs and SGC pairs. In some experiments, slices from were exposed to wide-field blue (λ=470nm) or amber (λ=589nm, experiments, slices from were exposed to wide-field blue (λ=470nm) or amber (λ=589nm, 10 =589nm, mW/mm^2^) illumination using Lambda DG-4 Plus (Sutter Instruments, Novato, CA) or Thorlabs 4-Wavelength High-Power LED Sources (Thorlabs, Newton, NJ).

### Immunohistochemistry

Following physiological recordings, slices were fixed in 0.1mM phosphate buffer containing 4% paraformaldehyde at 4°C. For *post-hoc* immunohistochemistry, slices were incubated overnight at 4°C with Alexa Fluor 594-conjugated streptavidin in 0.3% Triton X-100 and 2% normal goat serum containing PBS. Slices were mounted on glass slides using vectashield. Sections were visualized and imaged using a Zeiss LSM 510 confocal microscope with a 0.5 numerical aperture 20× objective.

### Morphometry

Stained hippocampal sections were visualized and imaged using a Nikon A1R laser scanning confocal microscope with a 20X, 0.5 NA objective lens. Cell reconstructions, from confocal image stacks, were performed using the directional kernels user-guided reconstruction algorithm in Neurolucida 360 (MBF Bioscience) followed by manual correction and validation in 3D. About 10-20 percent of each dendritic arbor was reconstructed manually (Gupta et al., 2020). Algorithms in Neurolucida Explorer (MBF Biosciences) were used to extract nonnominal or nonordinal somatodendritic morphological quantitative parameters for use in statistical comparisons and hierarchical cluster analysis. A total of 17 projection neurons in which the dendritic arbors were fully reconstructed were analyzed. A total of 42 somato-dendritic parameters (Defined in Gupta et al., 2020) from 17 morphologically reconstructed neurons, including both features measured in Neurolucida Explorer from the 3D reconstructions and parameters measured manually in 2D rendering (Neurolucida 360, MicroBrightfield) were analyzed.

### Quantification and Statistical Analysis

Data were tested for uniform distribution and each quantified variable was fit to the sum of two or more Gaussian functions and quality of fit determined using maximum likelihood analysis (MLA; v2test) to assess normal distribution of parameters within each cell type. Variables with a nonuniform distribution were used for subsequent cluster analysis. Hierarchical cluster analysis on principal components (HCPC) of morphological parameters was conducted using R version 3.5.0, using R package FactoMineR by an investigator (H. X.) blinded to cell types and age groups. Hierarchical clustering on the selected principal components was performed using Ward’s criterion with Euclidean distance to generate the dendrogram. The clustering partition was obtained from hierarchical clustering and improved with K-means method (Husson et al., 2010).

Individual spontaneous, miniature and evoked IPSCs were detected using custom software in IgorPro7.0 (WaveMetrics, Lake Oswego, OR) and different parameters were analyzed (Gupta et al., 2012). Events were visualized, and any “noise” that spuriously met trigger specifications was rejected. EPSCs were detected and analyzed using event detection in Clampfit. Kinetics and charge transfer were calculated from the averaged trace of all accepted sIPSC events. Rise time was measured as the time for amplitude to change from 20 to 80% of peak. Amplitude weighted τ_decay_ was calculated from a two-exponential fit to the IPSC decay. sIPSC charge transfer was calculated as the area under the curve of the baseline adjusted average sIPSC trace. Intrinsic properties were extracted by analyzing the IV traces in Clampfit. Input resistance was calculated in response to −100 pA current injection. Spike frequency adaptation ratio was calculated as interval between the first two spikes/interval between the two last spikes in response to +70 pA current injection. Sample sizes were not predetermined and conformed with those employed in the field. Significance was set to p<0.05, subject to appropriate Bonferroni correction. Statistical analysis was performed by paired and unpaired Student’s *t* test (Microsoft Excel 2007) or Two-way repeated measures ANOVA, One-way ANOVA or One-way ANOVA on Ranks followed by Pairwise Multiple Comparison using Holm-Sidak method or Dunn’s method (Sigma Plot 12.3) as appropriate. Data that passed normality test are shown as mean±SEM (standard error of the mean). Data that were not distributed normally are reported as median and interquartile range (IQR) as appropriate.

## Results

### Quantitative morphometry resolves GCs and SGCs in mice into distinct groups

Recordings were obtained from neurons in the inner molecular layer (IML) to target putative SGCs and granule cell layer to target GCs followed by *post hoc* recovery of cell morphologies from biocytin fills (Fig. 1A-E). Unsupervised clustering of morphometric data obtained from reconstruction of neurons in which dendritic arbors were fully recovered was used to verify whether the somato-dendritic parameters distinguishing GCs from SGCs in mice are similar to those characterized in rats (Williams et al., 2007; Gupta et al., 2020). Projection neurons, which included GCs and SGCs, were identified based on the presence of spiny dendrites in the molecular layer and axons with mossy fiber boutons entering the hilus and targeting CA3 (Gupta et al., 2020). Morphometric parameters of the cells reconstructed in 3D were obtained using automated algorithms in Neurolucida Explorer (Define in Gupta et al., 2020). Principal Component Analysis (PCA) of 42 morphometric parameters from 17 cells revealed that the first two principal components (PCs) explained more than 50% of the total variance in the data, while the first seven components explained 90% of the variance. Hierarchical cluster analysis on PCs (HCPC) on the first seven PCs suggested partitioning of cells into three clusters (Fig. 1F) as illustrated on the factor map produced by the first seven PCs (Fig. 1G). In spite of the exclusion of categorical variables such as soma location from HCPC, cells classified as GCs and SGCs by investigator (M.A.) segregated into two distinct clusters based on Dimension1 (Dim1). Interestingly, cells classified as GCs further segregated into two clusters (Clusters #1 and #2) based on Dim2. Variables more than 70% correlated with the 1st PC were identified as most responsible for PC formation. Evaluation of the quantitative variables underlying the clusters revealed that cluster #3, comprised of putative SGCs, was distinguished by a higher number of primary dendrites, greater dendritic angle, soma width, and cell surface area and lower dendritic complexity (Fig. 1H-J). While cells in Cluster #1 had fewer, typically one, primary dendrite, lower dendritic angle, and higher dendritic complexity (Fig. 1H-J), Cluster #2 segregated based on the presence of more fourth order segments, ends and intersections. Interestingly, dendritic angle was different between groups, increasing from Cluster #1 to Cluster #3 (Fig 1H). The number of primary dendrites in Cluster #3 (putative SGCs) was significantly higher than in both Clusters #1 and #2 (Fig 1I). Similarly, Cluster #3 (putative SGCs) had the least dendritic complexity which was significantly lower than in Cluster #1 while differences in dendritic complexity between Clusters #1 and #2 failed to reach statistical significance (Fig 1J).

**Figure 1:**
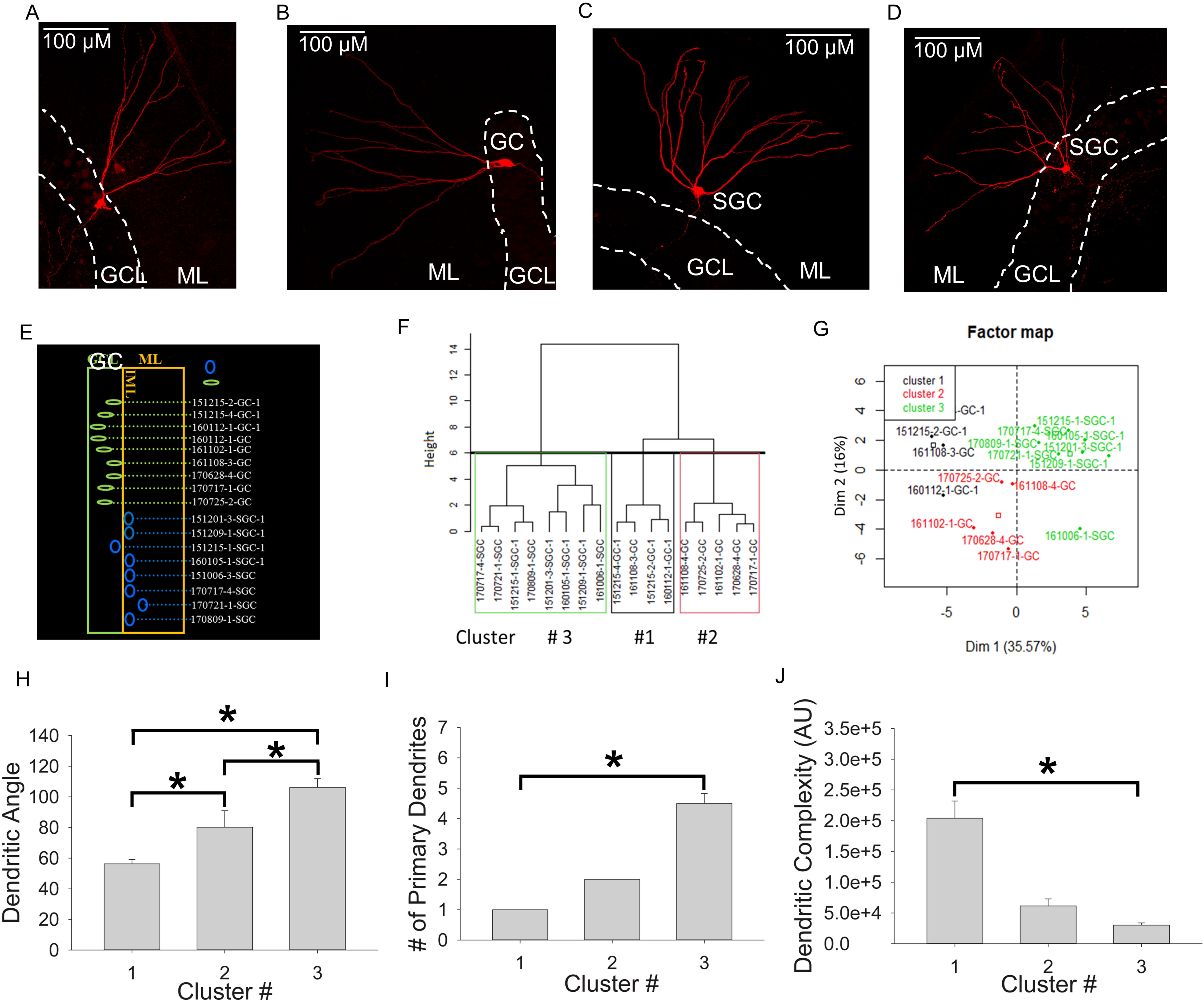
GCs and SGCs cluster into distinct subtypes on the basis of morphology. A-D. Example images of GCs (A-B) and SGCs (C-D) illustrate the somatic location and dendritic spread. Note that reconstruction of neuron in A is presented in Fig 2A. Scale bar: 100µm; E. Schematic of the somatic location of the cells included in the cluster analysis. The GCL is illustrated as a green box and IML as an orange box with somata illustrated in blue and GCs in green. All images were obtained a 20X magnification. GCL: Granule Cell Layer, IML: Inner Molecular Layer. F. Dendrogram generated by Hierarchical Clustering on Principal Components (HCPC) based on 42 morphometric parameters (Defined in Gupta et al., 2020) using Ward’s method with Euclidean distance suggests three putative clusters. G. Factor map base on first 7 PCs. H-J. Summary data of dendritic angle (H), number of primary dendrites (I) and Dendritic Complexity (J) for cells in the three clusters. Data are presented as mean±sem. * indicates p<0.05 by One-way ANOVA followed by pairwise comparison with Holm-Sidak method for H and p<0.05 by One-way ANOVA on Ranks followed by pairwise comparison with Dunn’s method for I and J where the data failed Shapiro-Wilk Normality Test. A. U: Arbitrary Units.

Morphometric parameters from putative GCs (Clusters #1 and #2) were pooled together and compared with Cluster #3 (putative SGCs) for further analysis. Like GCs, SGC axons extended to CA3 and showed mossy fiber terminals in the hilus (Fig. 2A-B), consistent with features of a projection neuron. Similar to findings in rats (Gupta et al., 2020), GCs and SGCs differed in dendritic angle, number of primary dendrites, soma aspect ratio and dendritic complexity (Fig. 2C-F, Table 1). In contrast, certain global parameters such as total number of dendritic ends and total length of dendrites were not different between the cell types (Fig. 2G-H, Table 1). Together these data provide a comprehensive morphometric comparison of GCs and SGCs in mice. Our results confirm that the presence of 3 or more primary dendrites and wide dendritic angle reliably distinguishes SGCs from GCs as reported in recent studies (Save et al., 2018; Gupta et al., 2020; Rovira-Esteban et al., 2020).

**Figure 2.**
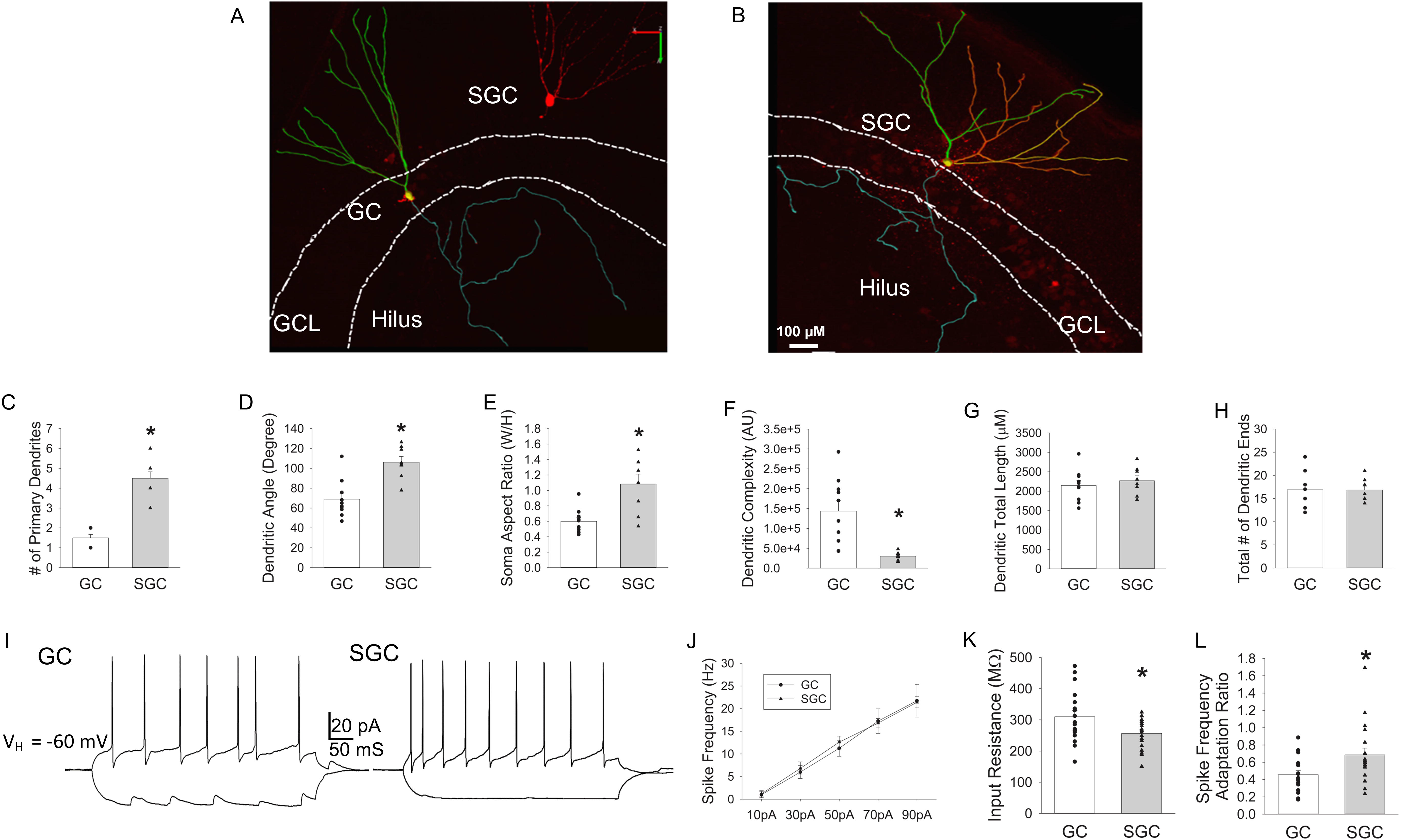
Morphological and physiological characteristics of GCs and SGCs in mice. A-B. Overlay of confocal image and Neurolucida 360 reconstruction of a biocytin filled GC (A) and SGC (B) show dendritic arbors and axon projection towards CA3. Cyan: axons; Orange, Green, Yellow: dendritic trees. C-H. Summary plots of morphological parameters including number of primary dendrites (C), dendritic angle (D), soma aspect ratio (E), dendritic complexity (F), dendritic total length (G) and total number of dendritic ends (H) between GCs (n=10) and SGCs (n=8). I. Voltage traces in response to +70 pA and −100 pA current injections in a GC (left) and SGC (right) illustrate firing pattern and passive properties. J. Summary plot of firing frequency in response to increasing current injections in GCs (n=8) and SGCs (n=10). K-L. Summary histogram of input resistance (measured from response to −100 pA), (K) and spike frequency adaptation ratio (L) in GCs (n=18) and SGCs (n=19). Data presented as mean±sem. * indicates p<0.05 by independent t-test.

**Table 1:** Summary of morphometric and physiology data in GC and SGCs.

| Fig # | Parameter (unit) | Cluster 1<br>(5 cells/ 8 mice) | Cluster 2<br>(4 cells /8 mice) | Cluster 3<br>(8 cells/ 8 mice) | F and P Value |
| --- | --- | --- | --- | --- | --- |
| 1H | Dendritic Angle (Degree) | 56.28±2.84 | 80.21±10.82 | 105.12±6.17 | F(2,14)=16.13<br>p <0.001 by One Way ANOVA<br>Clu1 v Clu2 =0.038<br>Clu1 v Clu3<0.001<br>Clu2 v Clu3=0.032 |
| 1I | # of Primary Dendrites (Count) | 1.00±0.00 | 2.00±0.00 | 4.60±0.40 | <0.001 by Kruskal-Wallis ANOVA on Ranks<br>Clu1 v Clu2 =0.038<br>Clu1 v Clu3<0.001<br>Clu2 v Clu3=0.032 |
| 1J | Dendritic Complexity (A.U.) | 204147±27920 | 61393±11452 | 30435±4609 | 0.002 by Kruskal-Wallis ANOVA on Ranks<br>Clu1 v Clu2 >0.05<br>Clu1 v Clu3<0.05<br>Clu2 v Clu3>0.05 |
| Fig # | Parameter (unit) | GC |  | SGC | P Value by Independent Student's t-test |
| 2C | # of Primary Dendrites (Count) | 1.50±0.17 |  | 4.5±0.33 | 1.90E-07 |
| 2D | Dendritic Angle (Degree) | 69.00±5.98 |  | 106.23±5.71 | 0.0004 |
| 2E | Soma Aspect Ratio | 0.60±0.05 |  | 1.08±0.13 | 0.001 |
| 2F | Dendritic Complexity (A.U.) | 143653.30±26506.76 |  | 30290.86±3515.42 | 0.002 |
| 2G | Total Dendritic Length (µM) | 2147.37±128.20 |  | 2269.97±126.30 | 0.512 |
| 2H | Total Dendritic Ends (Count) | 16.90±1.13 |  | 16.87±0.81 | 0.986 |
| 2K | Input Resistance (MΩ) | 309.89±19.58 |  | 229.33±29 | 0.031 |
| 2L | Spike Frequency Adaptation Ratio | 0.45±0.05 |  | 0.68±0.07 | 0.023 |

### SGCs differ from GCs in intrinsic physiology and frequency of basal synaptic inhibition

Examination of active and passive responses to graded current injections revealed no difference in firing frequency between the two cell types (Fig. 2I-K). However, input resistance was significantly lower in SGCs (Fig. 2K). Spike frequency adaptation ratio (ratio of inter-spike interval between first 2 and last 2 spikes during a +70 pA current injection) in SGCs was lower than in GCs (Fig. 2L). These findings are consistent with differences in intrinsic properties identified between the SGCs and GCs (Williams et al., 2007; Gupta et al., 2012; Save et al., 2018).

Spontaneous inhibitory postsynaptic currents (sIPSCs), recorded from holding potential of −60 mV in the presence of glutamate blockers, were more frequent in SGCs (Fig. 3A, C: sIPSC frequency in Hz, GCs: 8.38±1.14, N=18/12 mice, SGCs: 13.48±1.07, N=19/ 13 mice, p=0.003, t-test). However, sIPSCs amplitude (Fig. 3D: sIPSC amplitude in pA, GCs: 42.98±3.35, N=18, SGCs: 41.52±3.01, N=19, p=0.75, t-test), amplitude weighted decay time constant (τ_decay_-WT in ms, GCs: 3.31±0.22, N=18, SGCs: 3.2±0.2, N=19, p=0.72, t-test) and 20-80% rise times (in ms, GCs: 0.22±0.01, N=18, SGCs: 0.20±0.0, N=19, p=0.13, t-test) were not different between GCs and SGCs (Fig. 3B, D-F). These findings demonstrate that SGCs in mice show the same structural and functional distinctions from GCs identified in prior studies in rats (Gupta et al., 2012; Gupta et al., 2020). These data validate the morphological criteria which we used to distinguish GCs and SGCs in the subsequent experiments and indicate differences in inhibitory inputs between the cell types.

**Figure 3.**
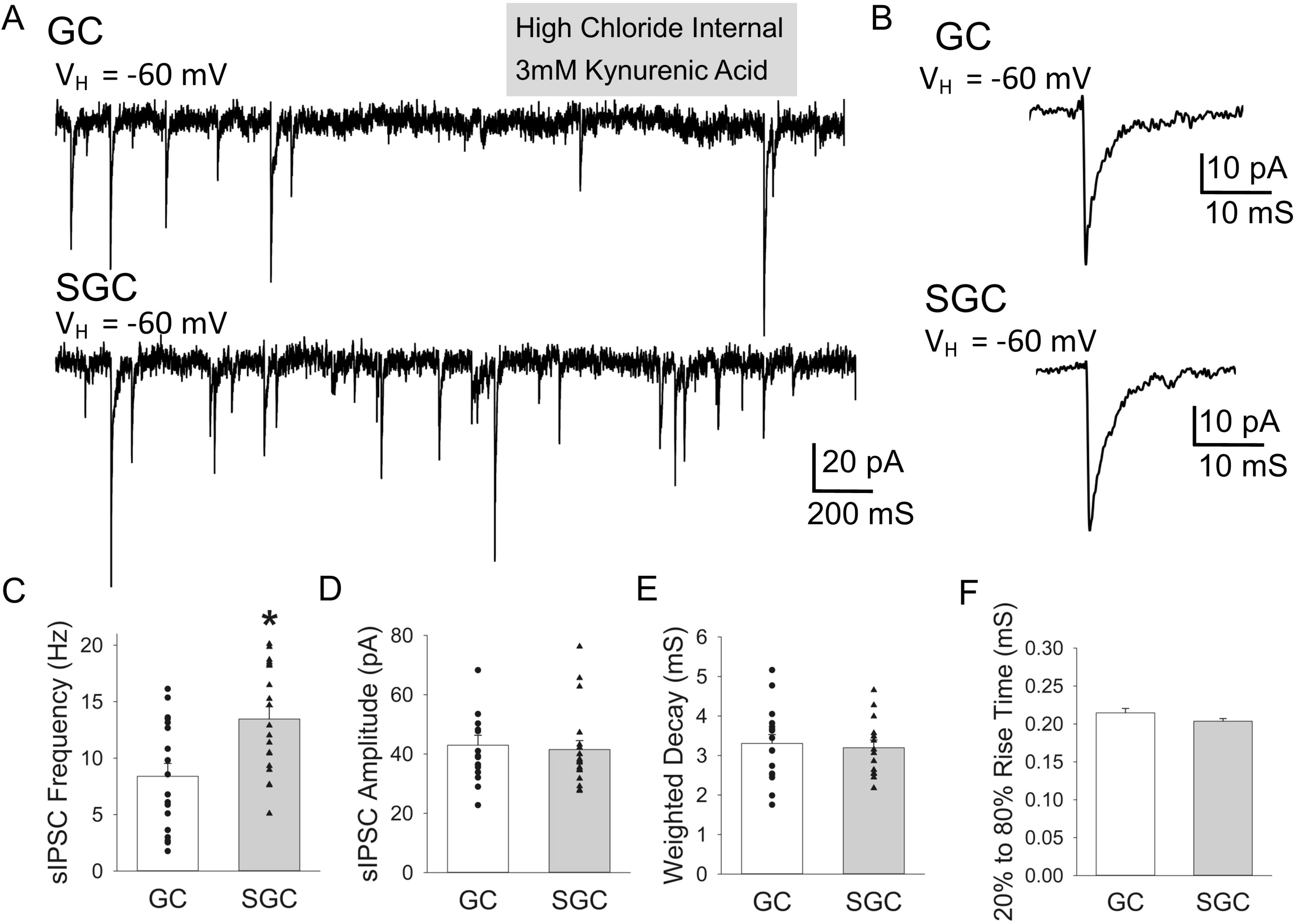
SGCs receive more spontaneous inhibitory synaptic events than GCs. A. Representative current traces from a holding potential of −60 mV illustrate sIPSCs in a GC (above) and SGC (below). Text in gray box indicates recording condition. B. Example of average sIPSC trace from a GC (above) and SGC (below). C-F. Summary histogram of sIPSC frequency (C), amplitude (D), amplitude weighted decay time constant (E) and 20-80% rise time (F) in GCs (n=18) and SGCs (n=19). Data presented as mean±sem. * indicates p<0.05 by independent t-test.

### Limited role of parvalbumin interneurons in steady-state synaptic inhibition of GCs and SGCs in slices

To determine whether differential inputs from parvalbumin expressing interneurons (PV-INs), contribute to the higher action potential-dependent sIPSCs frequency in SGCs we examined if optical suppression of PV-INs in mice expressing the inhibitory opsin, halorhodropsin (NpHR3), under control of the PV promotor reduced sIPSC frequency. Unlike archaerhodopsin and chloride conducting channelrhodopsins which can depolarize terminals and induce synaptic release, NpHR3 has been shown to hyperpolarize neurons without triggering transmitter release making it the ideal choice to examine synaptic release (Mahn et al., 2016). Validity of the PV-Cre mouse line was confirmed by colocalization of parvalbumin and YFP using immunostaining (not shown) and by fast, non-adapting firing in recordings from YFP positive neurons (Fig. 4A). Since activation of NpHR3 can reduce the chloride gradient (Alfonsa et al., 2015), light exposure was restricted to 2s periods. Recordings from YFP labeled neurons established that the 2s amber light illumination reliably and reversibly hyperpolarized YFP positive fast spiking neurons (n>10 cells) and suppressed action potential firing (Fig. 4A). Spontaneous IPSCs were recorded from GCs and SGCs for a total of seven minutes: Recordings started with a two-minute baseline followed by five one-minute cycles with a 2s exposure to amber light at the beginning of each cycle followed by time for recovery of the chloride gradient between light exposure (Fig. 4B). Spontaneous IPSC parameters measured during the 2s prior to light exposure were averaged and compared to the parameters averaged over the periods during the five cycles of light exposure. Contrary to expectations, optical suppression of PV-INs failed to reduce both frequency (in Hz, GC no-light: 14.19±2.5; GC Amber light: 13.6±2.4, N=5/3 mice, p=0.61, paired t-test) and amplitude (in pA, GC no-light: 46.65±9.62; GC Amber light: 45.13±5.94, N=5/3 mice, p=0.42, paired t-test) of GC sIPSCs (Fig. 4 C-E). Similarly, optical suppression of PV-IN firing failed to reduce sIPSC frequency (in Hz, SGC no-light: 15.62±2.6; SGC Amber light: 14.69.6±2.0, N=5/3 mice, p=0.79, paired t-test) and amplitude (in Hz, GC no-light: 43.18±5.1; GC Amber light: 40.95±3.2, N=5/4 mice, p=0.42, paired t-test) in SGCs as well (Fig. 4C-E). These data raise a surprising possibility that PV-INs may not contribute to basal sIPSCs in either GCs or SGCs in acute slice preparations.

**Figure 4.**
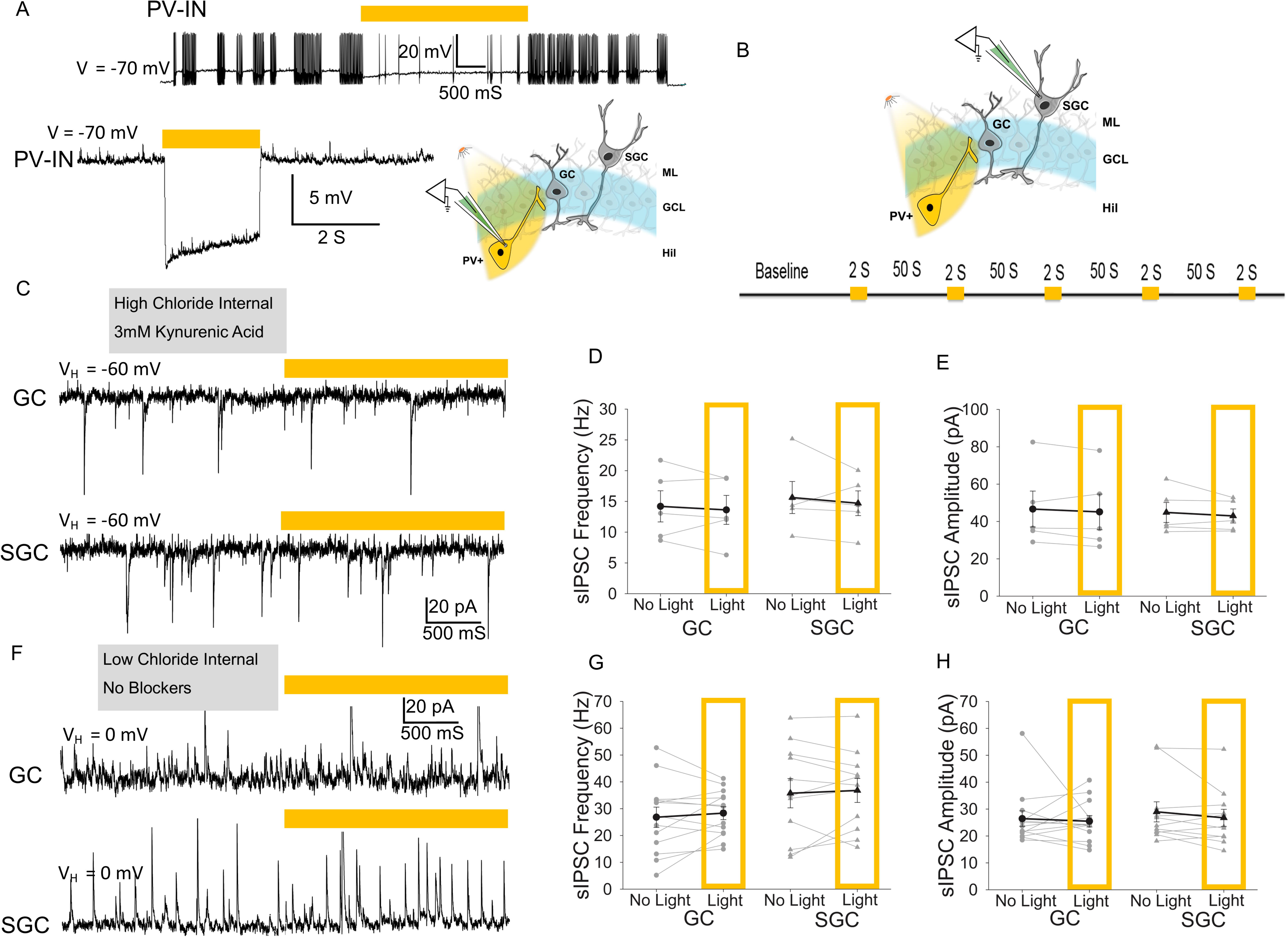
Optical suppression of PV-INs fails to reduce sIPSC frequency in both GCs and SGCs. A. Representative membrane voltage trace (above) from a holding potential of −70 mV in a YFP-positive PV interneuron expressing halorhodopsin in the granule cell-hilar border illustrates the high frequency firing during depolarizing current injection (100 pA for 10 sec) and the ability of amber light (bar above) to suppress firing. Representative trace from 19 trials in 4 cells. Membrane voltage trace from an NpHR3-YFP expressing PV-IN shows rapid and reversible hyperpolarization (below) during the 2s amber light stimulation. Schematic to the right illustrates recording condition and optical stimulation. B. Schematic of the experimental paradigm illustrates amber light stimulation of NpHR3-YFP expressing PV-IN while recording from GC/SGC (above) and the protocol for light OFF and light ON condition below. C. Current traces from a GC (above) and SGC (below) recorded from a holding potential of −60 mV using high-chloride internal shows sIPSCs as inward currents before and during NpHR3 activation (amber bar) in PV-INs. IPSCs were isolated in kynurenic acid (3 mM). D-E. Plots summarize effect of optical suppression of PV-IN on sIPSC frequency (D) and amplitude (E) (n=5 GCs and SGC). F. Representative traces illustrate sIPSCs in a GC (above) and SGC (below) before and during activation amber light exposure (amber bar). Recordings were obtained using a low-chloride internal from a holding potential of 0 mV without synaptic blockers. G-H. Summary plots of effect amber light on sIPSC frequency (G) and amplitude (H) in GCs (n=13) and SGCs (n=11). Data presented as mean±sem. Text in gray box indicates recording condition.

Given the unexpected outcome, and since the optogenetic system was functionally validated (Fig. 4A), we considered whether use of glutamate receptor antagonist to isolate IPSCs may have reduced PV-IN firing and underestimated their contribution to sIPSCs. To address this possibility, IPSCs were recorded as outward currents from cells held at 0 mV (reversal for glutamatergic currents) using a low chloride internal. As expected, sIPSC frequency recorded in the absence of glutamate blockers was higher (Fig 4F-H) than that recorded in glutamate receptor blockers (Fig 3 and 4C-D) in both cell types. Nevertheless, sIPSC frequency and amplitude in GCs were not reduced by amber light suppression of PV-INs (Fig. 4F-H: frequency in Hz, no-light: 26.80±3.78, light: 28.32±2.41, N=13/ 8 mice, p=0.49, paired t-test; amplitude in pA, no-light: 26.43±2.88, light: 25.46±2.10, N=13/ 8 mice, p=0.76, paired t-test). Similarly, sIPSC parameters in SGCs were also unaltered during amber light exposure (Fig. 4F-H: frequency in Hz, no-light: 35.77±5.44, light: 36.81±4.48, N=11/ 8 mice, p=0.66, paired t-test; amplitude in pA, no-light: 28.96±3.72, light: 26.74±3.17, N=11/ 8 mice, p=0.25, paired t-test). Together, these data obtained under two different recording conditions support the surprising conclusion that PV-INs contribute minimally to basal sIPSC frequency in both dentate projection neuron subtypes.

While unlikely, a potential reason for the above outcome is that a majority of the sIPSCs in our recordings were action potential independent miniature events, which would not be modified by suppressing PV-IN firing. To exclude this possibility, we tested the ability of the sodium channel antagonist tetrodotoxin (TTX, 1μM) to reduce frequency of IPSCs indicating the presence of action potential driven events. As expected based on prior studies (Otis et al., 1991; Goswami et al., 2012), TTX significantly reduced IPSC frequency in both GCs (frequency in Hz, before ACSF: 11.58±1.58, TTX: 1.4±0.44 N=7, p=0.0003, paired t-test) and SGCs (frequency in Hz, before ACSF: 10.83±2.09, TTX: 1.96±0.9 N=5, p=0.001, paired t-test). TTX also reduced IPSC amplitude in both cell types (not shown). Overall, the ability of TTX to suppress IPSC frequency indicates that majority of sIPSCs in both SGCs and GCs are driven by interneuron firing. Taken together, these data support the conclusion that PV-INs do not contribute significantly to action potential driven basal sIPSCs in both GCs and SGCs in slice preparations.

### Activation of parvalbumin interneurons drives synaptic inhibition in both GCs and SGC

Since sIPSCs represent synaptic inhibition in the absence of afferent inputs, we examined whether direct optical activation of PV-IN evoked IPSCs (oeIPSCs) in GCs and SGCs in slices from mice expressing channelrhodopsin (ChR2) in PV-INs. Following baseline recordings of sIPSCs in glutamate antagonist without light, recordings were continued during optical activation of ChR2 in PV-IN using blue light (λ=473nm, 10 s). Optical activation of PV-INs elicited a significant increase in IPSC frequency in GCs (Fig. 5A-C; frequency in Hz, no-light: 8.45±1.31, light: 52.11±5.92, N=14/ 8 mice, p=2E-06, paired t-test) confirming the robust PV-IN input to GCs. PV-IN activation increased IPSC frequency in SGCs as well (Fig. 5A-C: frequency in Hz, no-light: 13.68±1.20, light: 59.49±4.62, N=14/ 9 mice, p=2E-07, paired t-test), demonstrating functional PV-IN innervation of SGCs. However, IPSC peak amplitude was not increased by optical activation of PV-INs in either GCs or SGCs (Fig. 5D-F). These data demonstrate that SGCs, like GCs, receive PV-IN inputs and that direct activation of PV-INs leads to an increase in synaptic inhibitory events in both GCs and SGCs. Since PV-INs are known to be recruited during PP activation and support feedforward and feedback GC inhibition (Ewell and Jones, 2010), we examined whether recruitment of inhibition by afferent stimulation differed between the cell types. Single PP stimulation at increasing intensities evoked IPSCs in both SGCs and GCs. The peak amplitude of afferent evoked IPSCs was not different between cell types (Fig. 5E-F, N=12 cells each from 8 mice, F(1,10)=0.874, p=0.37 by Two-Way Repeated Measures ANOVA), demonstrating that both GCs and SGCs receive similar early feedforward/feedback inhibition upon single PP stimulation.

**Figure 5.**
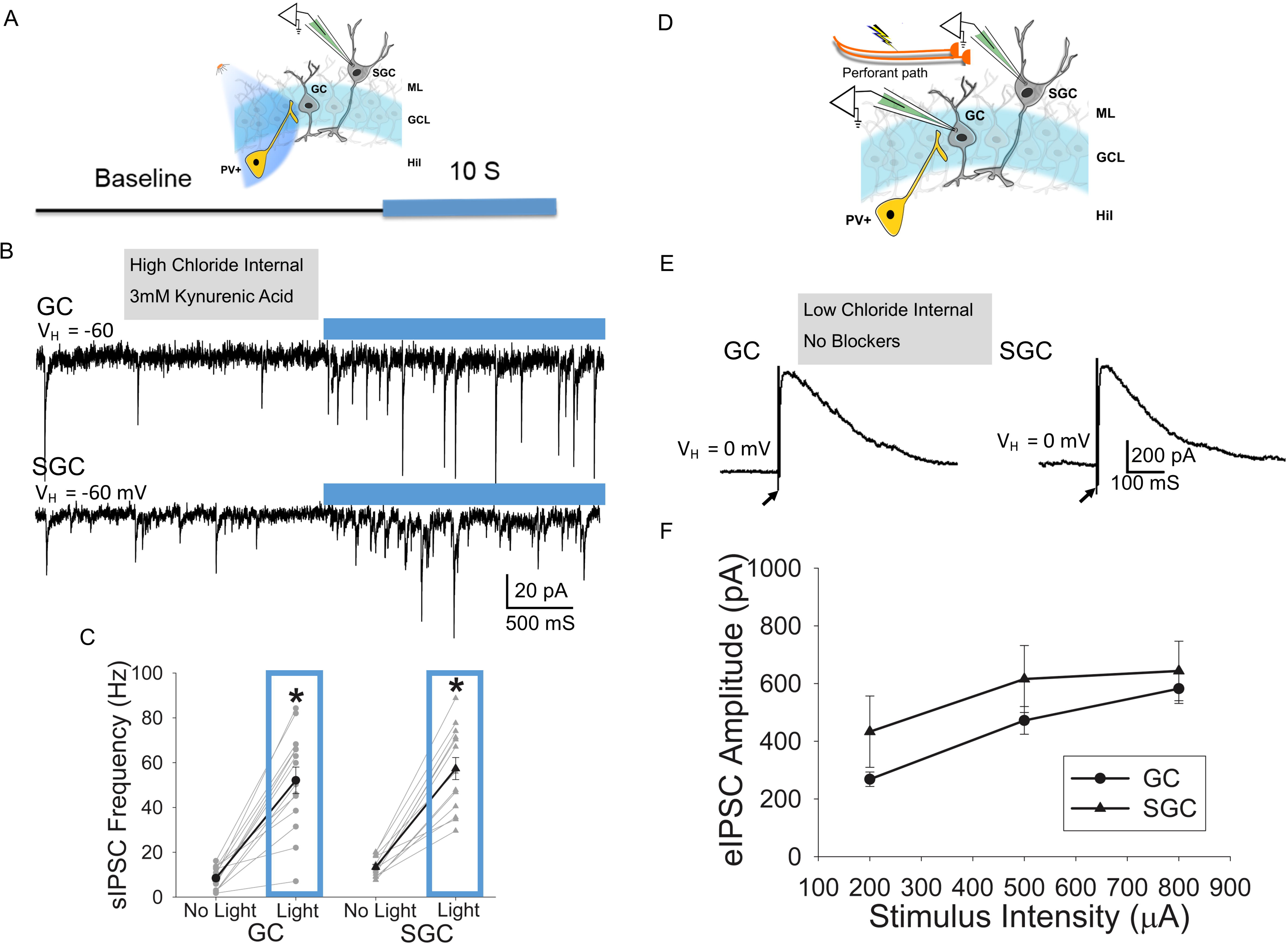
Activation of PV-INs and afferent stimulation evoke inhibitory synaptic currents in both dentate projection neuron subtypes. A. Schematic illustrates blue light stimulation of ChR2-YFP expressing PV-IN while recording from GC/SGC (above) and light OFF and light ON protocol below. B) Example recordings from a GC (above) and SGC (below) illustrate sIPSCs before and during optical activation of PV-INs. Cells were held at −60 mV and kynurenic acid (3 mM) was used to isolate IPSCs. C. Summary plots of sIPSC frequency before and during blue light exposure in GCs (n=14) and SGCs (n=14). D. Schematic of the experimental paradigm illustrates PP stimulation while recording from GC/SGC. E. Perforant path evoked IPSC in GC and SGC recorded as outward currents using a low-chloride internal and a holding potential of 0 mV. F. Summary plot of eIPSC peak amplitude in GCs and SGCs in response to stimulation of the PP at increasing current intensities. Data presented as mean±sem. * indicates p<0.05 by paired t-test (in panel C) or unpaired t-test (in panel D). Text in gray box indicates recording condition.

### Afferent stimulation drives greater enhancement of sustained feedback inhibition in granule cells

A functional characteristic of SGCs, not observed in GCs, is their unique ability for sustained firing in response to paired afferent stimuli in rat hippocampal slices, which has been proposed to drive feedback inhibition and sculpt pattern separation (Larimer and Strowbridge, 2010). We examined the reliability and reproducibility of SGC sustained firing in SGCs in mice. As is typical for GCs, a single PP stimulus evoked a single action potential in GCs (3/3 cells tested) as well as in SGCs (4/4 cells tested) but failed to elicit sustained spiking (not shown). However, when the PP was stimulated with a pair of suprathreshold stimuli at 10 ms interval, consistent with theta frequency inputs (Larimer and Strowbridge, 2010; Braganza et al., 2020), a subset of SGCs showed prolonged spiking activity. Sustained firing was observed in 51.51% of the trials (Fig 6A-B, 17/33 of trials in 2/4 SGCs). The sustained firing was initiated between 200ms to 2s after the paired stimulation, although the latency to sustained firing was over 6s in a few trials indicating that the sustained firing was mediated by network effects (Fig. 6C, median 1.14s IQR: 0.61-1.9s). This evoked sustained firing in SGC lasted several seconds, at times until the end of the recording 20s after stimulation (Fig. 6D). On average, the sustained firing lasted 15s, which is likely an underestimate due to the termination of recordings. None of the GCs tested exhibited firing beyond the early evoked firing in any of the trials (0/12 trials, 3 GCs).

**Figure 6.**
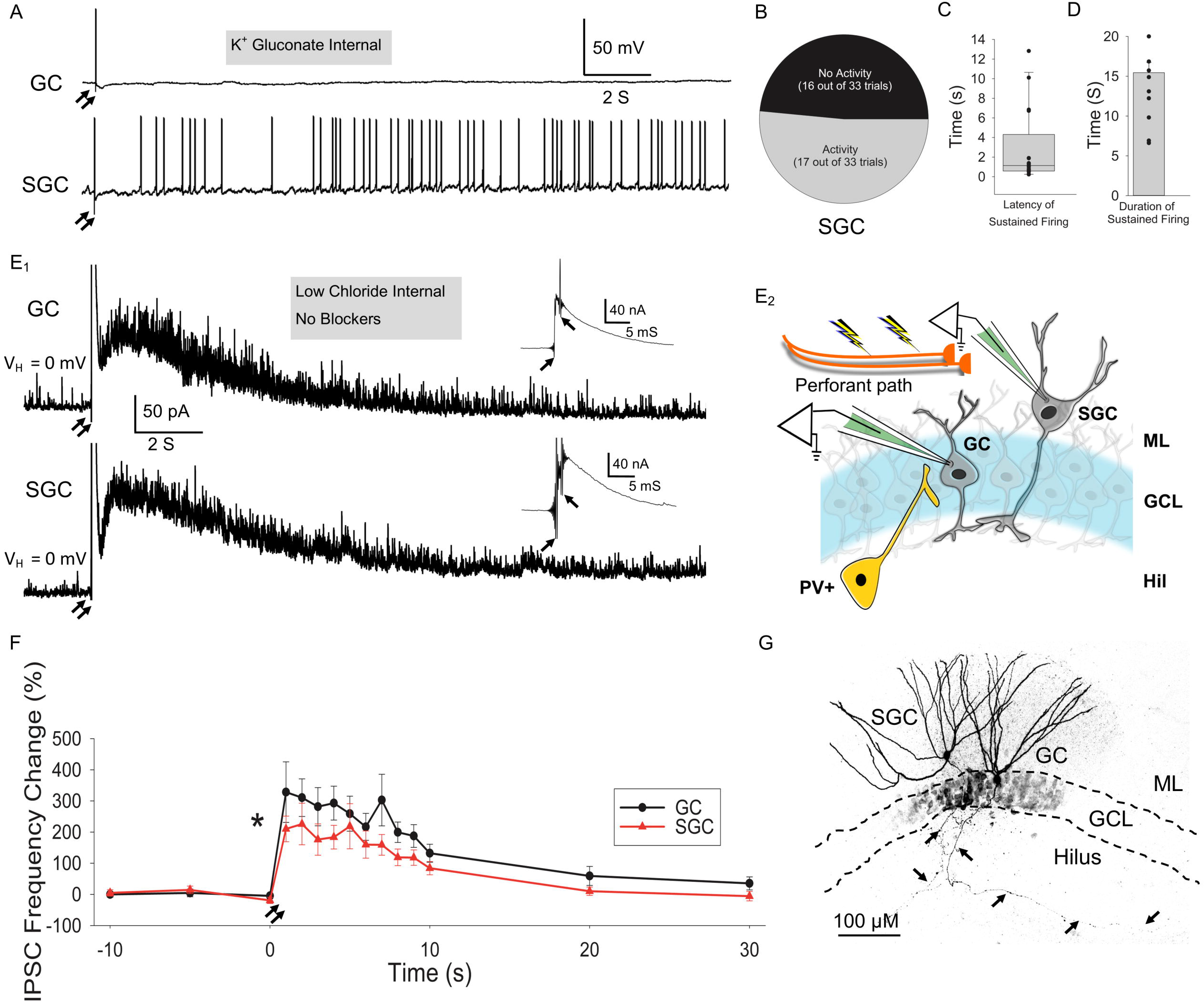
Afferent evoked prolonged SGC firing is associated with sustained increase in sIPSC frequency in both GC and SGC. A. Membrane voltage traces in a GC (above) and SGC (below) evoked by a pair of suprathreshold stimuli to the PP (denoted by 2 arrows). Note prolonged firing in the SGC while none of the GCs (n=3 cells, 12 trails) showed more than 1-3 spikes. B. Plot shows the proportion of trials that evoked sustained firing in SGCs (n= 4 cells, 33 trials). C-D. Plots illustrate the distribution of latency from afferent stimulation to the start of sustained firing (C) and duration of sustained firing in SGCs recorded for a maximum of 20s after stimulation. E. Schematic of paired GC-SGC recordings in response to paired PP stimulation (E_2_). Current traces of simultaneously recorded GC-SGC pair illustrate the sustained increase in IPSC frequency compared to pre-stimulus baseline (E_1_). The eIPSC peak is truncated to better illustrate the sustained change in IPSCs over 20s. E_1_ Insets: traces at a different scale illustrate the peak eIPSC amplitude. F. Summary plot of sIPSC frequency in GCs and SGCs (n= 12 each) prior to stimulation (binned over 5s) and evoked sustained IPSC frequency (1s bins for the first 10s and 5sec bins at 20 and 30s) for success rate in induction of prolonged spiking activity in GCs (12 trials, N=3) and SGCs (33 trials, N=5). G. Maximum intensity projection of a confocal image stack of a GC-SGC pair filled during recordings (presented as grey scale and inverted) illustrate dendric arbors and hilar axon (arrows). GCL: granule cell layer; ML: molecular layer, GC: granule cell; SGC: semilunar granule cell. Data presented as median (interquartile range) for in C and mean±sem in D and F. * indicates p<0.05 for effect of cell-type (F(1,11)=4.9) by TW-RM ANOVA. Text in gray box indicates recording condition.

Sustained firing in SGCs has been correlated with hilar mossy cell and interneuron firing as well as reduced GC excitability (Larimer and Strowbridge, 2010). However, whether GCs receive enhanced synaptic inhibition during this period and whether SGCs undergo a parallel sustained increase in feedback synaptic inhibition is not known. Using simultaneous recordings in GC-SGC pairs (Fig 6E), we directly compared recruitment of polysynaptic sustained inhibition in GCs and SGCs following paired PP stimulation. Representative image of cells filled during paired recordings show the GC with soma located in the cell layer and SGC somata in the inner molecular layer (Fig. 6G). Paired PP stimuli elicited a large early eIPSC peak (Fig. 6E_1_ insets) followed by a prolonged barrage of “evoked-sustained” IPSCs (esIPSCs) lasting over 20s in both GCs and SGCs (Fig. 6E_1_). To better characterize the evoked sustained activity, we binned IPSCs into 5s segments before the stimulus and 1s segments 500ms-10.5s after the stimulation, with additional 5s segments 15-20s and 25-30s post-stimulus (Fig. 6F). The 500ms immediately following stimulation was not included in the analysis in order to avoid the period of maximum evoked IPSC conductance which included summated events and obscured individual events. The peak IPSC frequency following paired PP stimulation (500-1500 ms) showed high variability and was not different between cell types (in Hz, GC: 64.7±4.3; SGC: 72.9±5.3, n=12 each p>0.05 by unpaired Student’s t-test). Although the sIPSC frequency prior to stimulation was stable within each cell, there was considerable cell-to-cell variability. To normalize for the variability and assess relative change in esIPSC frequency, the difference in IPSC frequency following stimulation was normalized to the pre-stimulus sIPSC frequency averaged over 3 × 5-sec pre-stimulus bins ((post-pre)/pre*100). Both GCs and SGCs showed a significant post-stimulus increase in esIPSC frequency lasting over 20s. In addition to the significant effect of time, the difference between cell types was statistically significant (Fig. 6F, F_(1,11)_=9.4, for effect of time and F_(1,11)_=4.9 for cell-type, p<0.05 by Two-way RM ANOVA). Pairwise comparison between cell types, revealed that the increase in esIPSC frequency normalized to pre-stimulus frequency was significantly greater in GCs than in SGCs. Both GCs and SGCs showed increase in esIPSC frequency for up to 10s post stimulus reaching similar maximum frequency and declining back to pre-stimulus levels by 20-30s. However, the relative increase in esIPSC frequency (normalized to pre-stimulus frequency) quantified over the first 500-1500ms (1s) after stimulation was significantly higher in GCs than in SGCs (p<0.05, by TW-RM ANOVA followed by Pairwise Multiple Comparison using Holm-Sidak method).

Interestingly, not all cells showed robust esIPSC barrages even within the same slice, suggesting cell specific recruitment of feedback inhibition. We found that 16.67% (2/12) GCs and 33.33% (4/12) SGCs failed to show esIPSC barrage, defined as doubling in basal frequency in the first 3 post-stimulus bins examined (3s). Additional recordings conducted in the presence of the GABA_A_ receptor antagonist, gabazine (10 µM) confirmed that, like eIPSC, esIPSCs were also eliminated by gabazine (not shown). These results demonstrate that paired PP stimulation results in prolonged changes to dentate inhibition and SGC excitability. Crucially, the data reveal that the relative increase in sustained feedback inhibition in SGCs is less than in GCs and suggest that in spite of their higher basal synaptic inhibitory tone, SGC show lower increase in inhibitory gating than GCs following activation of inputs.

### Role of PV-INs in sustained feedback inhibition of dentate projection neurons following paired afferent stimulation

Since PV-INs are known to mediate strong granule cell feedforward and feedback inhibition (Hefft et al., 2002), we examined whether they exhibit sustained firing in response to paired PP stimuli coinciding with increased esIPSCs in GCs. PV-INs in the hilar granule cell border were identified based on YFP labeling and confirmed by the high frequency, non-adapting firing (Fig. 7A_1_, insets). PV-IN voltage traces in responses to paired PP stimulation (Fig. 7A) are illustrated alongside the time course of esIPSC in a GC (Fig. 7D) for reference. A subset of PV-INs (4/11 cells, 36.4%) exhibited spontaneous action potential firing at rest under our recording conditions. Interestingly, only 54.5% (6/11 cells) PV-INs showed sustained increase in firing upon paired PP stimulation. An additional 18.2% (2/11 cells) PV-INs received increased EPSC drive during this period suggesting that they may be driven by SGC firing (Fig. 7B). However, 27.3% (3/11) PV-INs showed no apparent sustained change in EPSC frequency upon PP stimulation in spite of the early evoked firing. The prolonged PV-IN firing followed the initial stimulation with a median latency of 1.02s (IQR 0.62-2.03s) which was similar to the latency to evoked SGC firing (Fig. 7C). Since SGCs and not GCs show evoked sustained firing following paired perforant path stimulation, these data suggest that the sustained firing in SGCs likely drives PV-IN firing. These results raise the possibility that PV-IN firing contributes to a certain component of esIPSCs in GCs and SGCs.

**Figure 7.**
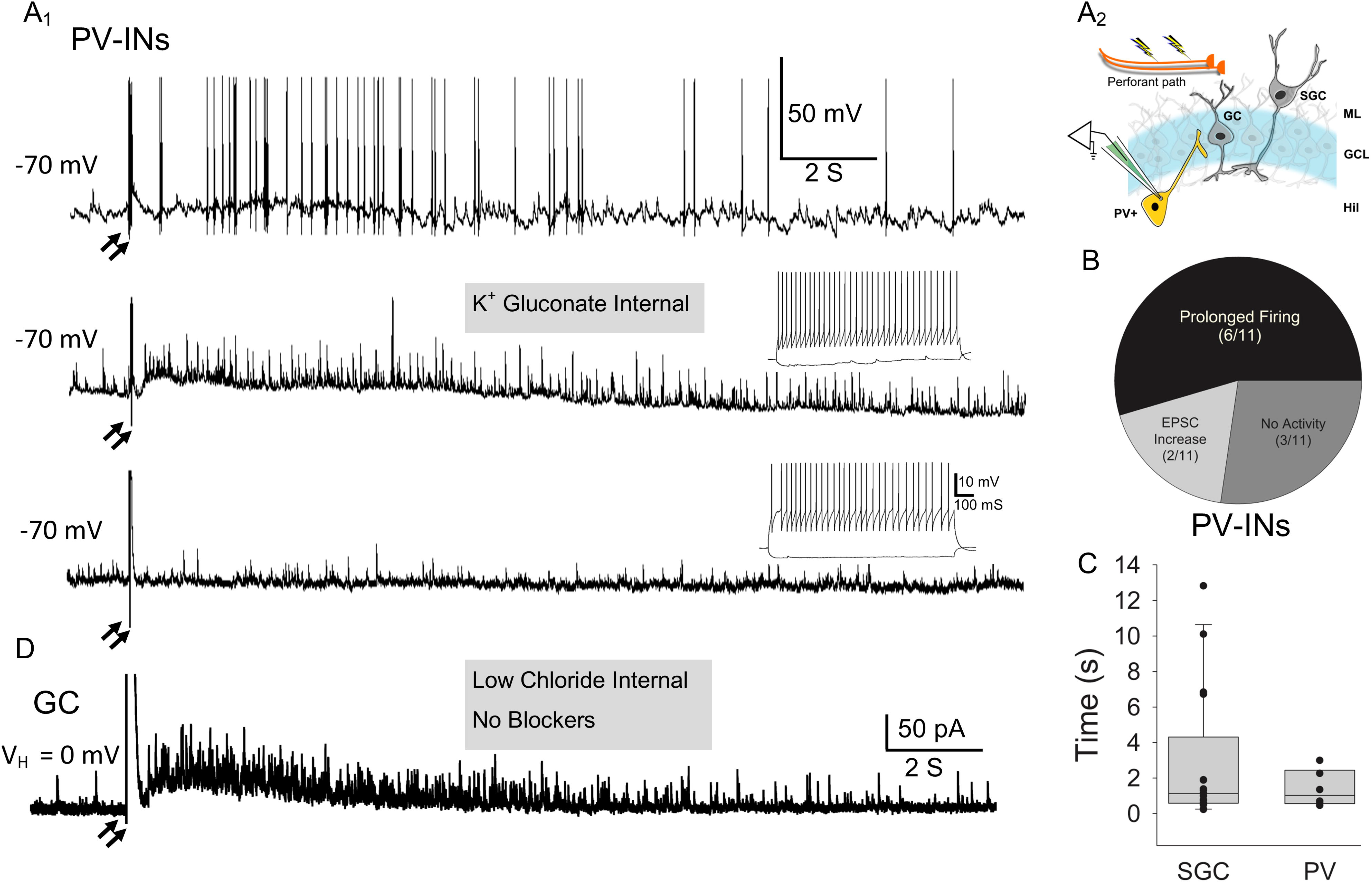
Increase in PV-IN excitability during evoked sustained sIPSCs in dentate projection neurons. A. Example traces show increased firing (above), enhanced EPSP barrages (middle) and unaltered EPSP frequency (below) in YFP-positive PV-INs (A_1_) in response to the paired PP stimulation paradigm (denoted by 2 arrows) illustrated in A_2_. Insets in middle and lower traces illustrate high frequency firing in recorded PV-IN. B. Pie chart summarizes the proportion of PV-INs that showed evoked sustained firing, EPSP increase or no change in response to stimulation (n= 11 cells, 33 trials). C. Plot illustrates the distribution of latency of enhanced PV-IN firing following paired PP stimulation. D. Current traces from a GC showing sustained IPSCs is included for comparison. Text in gray box indicates recording condition.

To verify whether halorhodopsin suppression could reduce PP-evoked PV-IN firing, we performed control recordings from YFP-labeled neurons in PV-NpHR3 mice. While optical suppression failed to fully eliminate firing of PV-INs in response to paired PP stimulation it consistently reduced firing without altering firing in hilar neurons lacking YFP expression (Fig. 8A-D). Additionally, we verified that optical suppression of PV-INs did not induce persistent firing in GC or eliminate sustained firing in SGCs (Fig. 8E).

**Figure 8:**
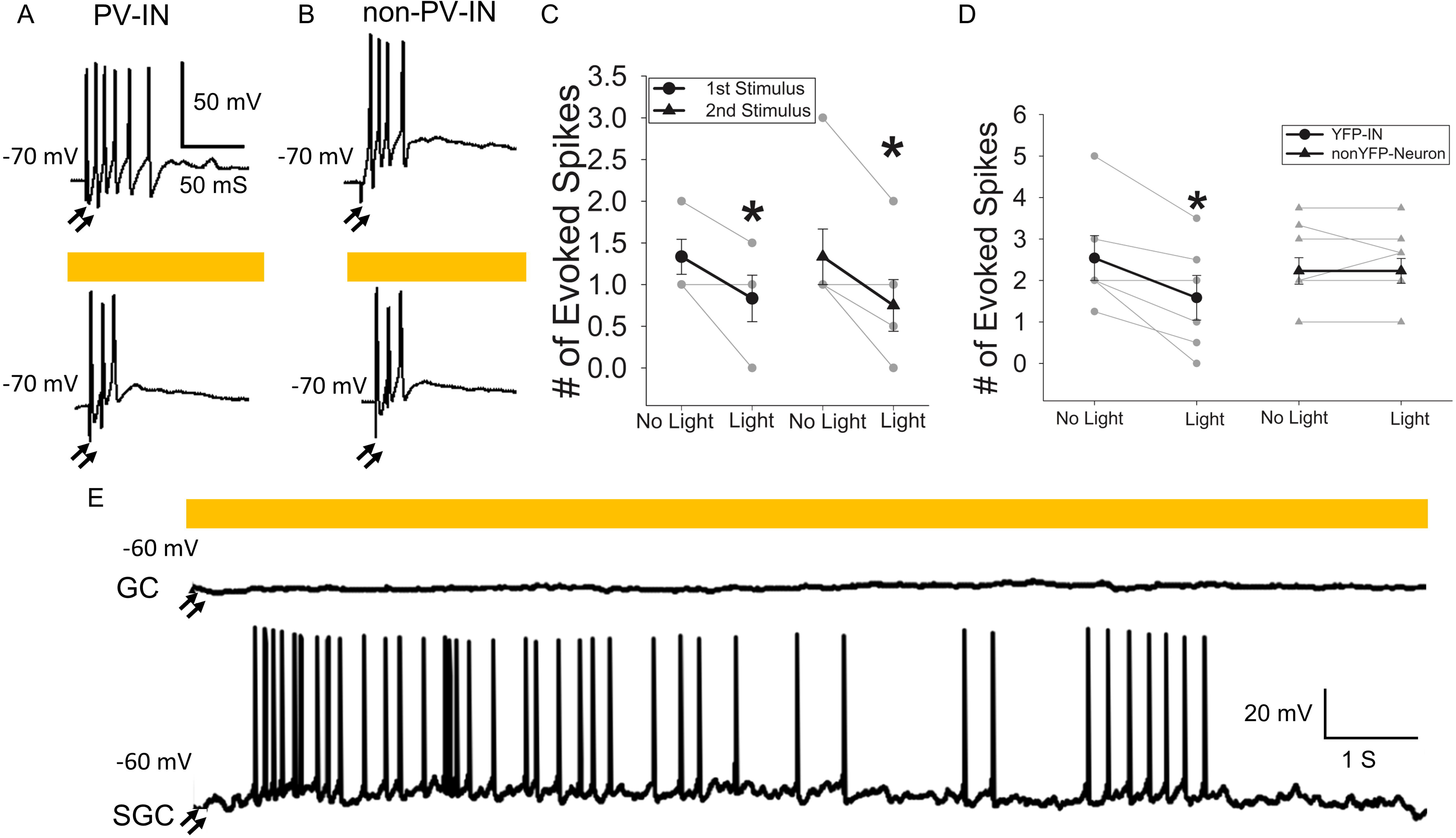
Cell specificity of halorhodopsin suppression of PV neuron firing. A. Representative membrane voltage traces from a YFP-positive PV interneuron expressing halorhodopsin in the granule cell-hilar border in response to a pair of suprathreshold perforant path stimuli (denoted by 2 arrows) at 10 ms interval illustrated the early evoked firing in PV neurons (above). Effect of amber light exposure on perforant path evoked PV-IN firing is illustrated (below). B. Example voltage traces from a YFP-negative hilar neuron in response to a pair of suprathreshold perforant path stimuli at 10 ms interval (above). Effect of amber light exposure on perforant path hilar neuron firing is illustrated (below). C. Summary data of number of spikes elicited following the first and second stimuli and the effect of halorhodopsin activation on perforant path evoked firing in PV-IN. D. Summary plot compares the effect of amber light on afferent evoke firing in halorhodopsin expressing PV-IN and YFP-negative hilar neurons. * Indicates p<0.05 by paired t-test. Data presented as mean±sem. E. Representative traces show lack of sustained firing in GCs and sustained firing in SGCs following paired suprathreshold perforant path stimulation. Text in gray box indicates recording condition.

To directly test the contribution of PV-INs to esIPSC barrages in GCs and SGCs, we recorded esIPSCs in putative GC/SGC pairs in PV-NpHR3 mice in the absence or presence of amber light (2s exposure starting 100ms after the onset of PP stimulation) (Fig. 9A-C). Light-on and light-off conditions randomized (Fig. 9, trials with red lined in Fig. 9D indicate light-on first recordings). Surprisingly, esIPSC frequency, normalized to basal sIPSC frequency prior to stimulation, was not reduced by optical suppression of PV-INs (Fig. 9D: esIPSC frequency as % of pre-stimulus frequency, no-light: 264.59±49.76, light: 225.85±43.08, N=8, p>0.05, by paired t-test). In contrast, recordings from SGCs showed a significant reduction in esIPSC frequency during suppression of PV-INs with amber-light (Fig. 8D: esIPSC frequency as % of pre-stimulus frequency, no-light: 209.76±57.92, light: 127.86±41.15, N=8, p= 0.015 by paired t-test). Thus, optical suppression of PV-INs selectively reduced esIPSC frequency in SGCs to approximately half of that observed in the absence of light demonstrating fundamental differences in inhibitory regulation of GCs and SGCs during evoked activity.

**Figure 9.**
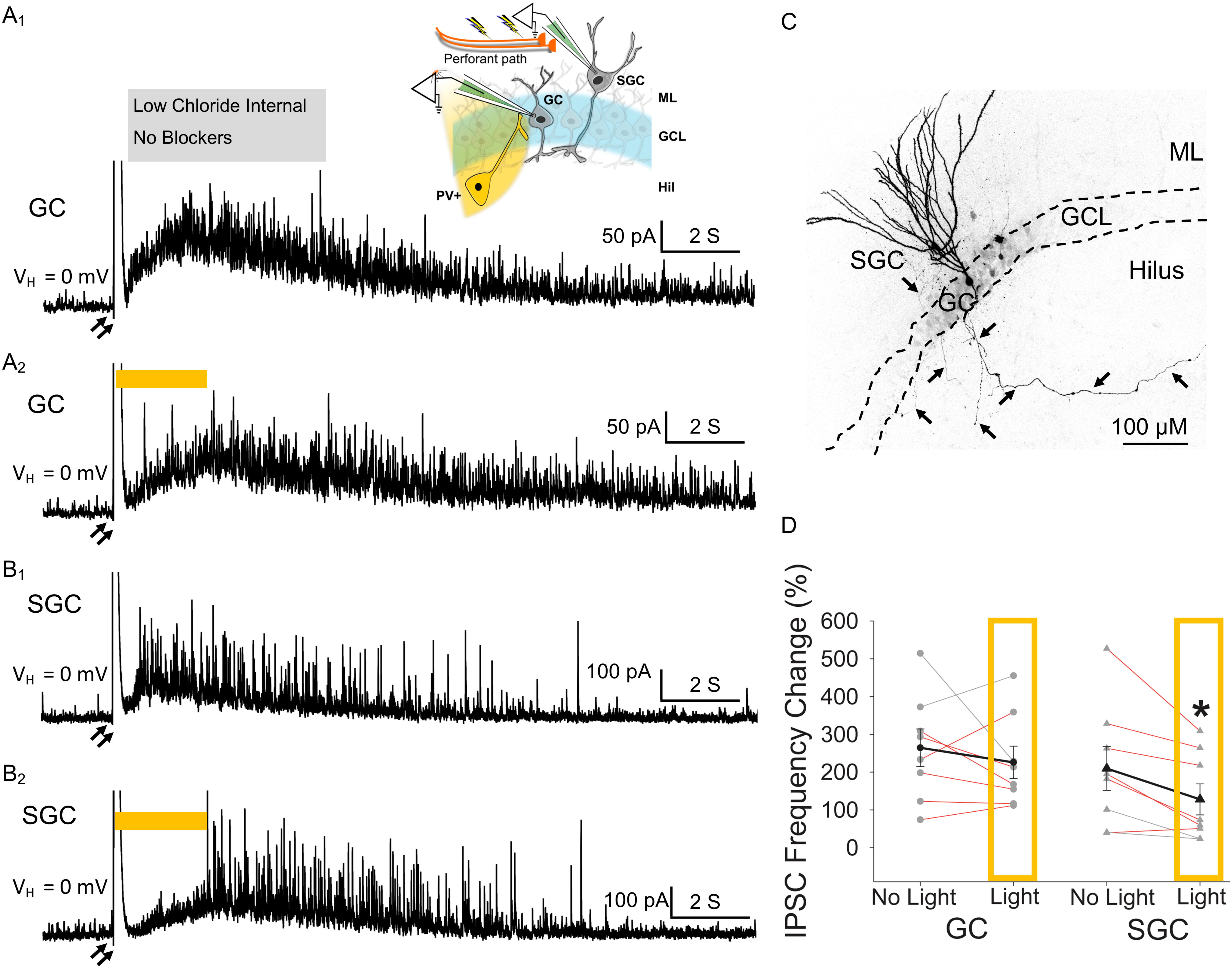
Optical suppression of PV-INs selectively reduces evoked sustained IPSCs in SGCs. A_1_. Schematic illustrating the simultaneous dual GC-SGC recordings in response to paired stimulation of the PP and amber light stimulation (above). GC current trace illustrates the sustained increase in IPSC frequency compared to pre-stimulus baseline (below). A_2_. GC trace shows that evoked sustained IPSCs in response to paired PP stimuli (denoted by 2 arrows) appears unchanged by amber light suppression (amber bar) of PV-INs. B_1_. SGC current trace illustrates the sustained increase in IPSC frequency compared to pre-stimulus baseline. B_2_. SGC trace shows that evoked sustained IPSCs in response to paired PP stimuli are suppressed by amber light. Note that the eIPSC peak is truncated to better illustrate the sustained change in IPSCs over 20s. C. Maximum intensity projection of a confocal image stack of a GC-SGC pair filled during recordings illustrated the soma location dendric arbors and hilar axon (arrows) of the cells. The image is presented as grey scale and inverted to better illustrate the axon. GCL: granule cell layer; ML: molecular layer, GC: granule cell; SGC: semilunar granule cell. D. Summary plots of effect of amber light suppression of PV-INs on evoked sustained IPSC frequency in GCs and SGCs (n=8 each). Red lines indicate trials with light-on first recordings. Data presented as mean±sem. * Indicates p<0.05 by paired t-test. Text in gray box indicates recording condition.

## Discussion

Here we identify that lateral sustained feedback inhibition, which has been proposed to shape dentate pattern separation and memory engram formation, is significantly lower in dentate SGCs than in GCs. We further show that parvalbumin interneurons selectively support activity-driven sustained lateral inhibition of SGCs, but not GCs. The difference in local circuit feedback inhibitory regulation of SGCs and GCs during afferent activation could shape their distinct role in dentate processing.

SGCs, originally defined by Ramón y Cajal (1995) are a sparse, functionally distinct subpopulation of dentate output neurons with CA3 projections (Williams et al., 2007; Larimer and Strowbridge, 2010; Gupta et al., 2012; Save et al., 2018). Temporally structured inhibition by diverse interneurons projecting across the somato-dendritic axis help maintain the sparse granule cell activity needed for pattern separation (Arima-Yoshida et al., 2011; Dengler and Coulter, 2016; Hainmueller and Bartos, 2020). Granule cells receive distinct phases of synaptic inhibition including steady-state spontaneous synaptic inhibition, afferent evoked mono-and disynaptic feedforward and feedback inhibition, as well as extrasynaptic inhibition (Soltesz et al., 1995; Kraushaar and Jonas, 2000; Stell et al., 2003). Additionally, afferent stimulation has been proposed to recruit sustained polysynaptic feedback GC inhibition (Larimer and Strowbridge, 2010). Focusing on synaptic inhibition, we demonstrate three functionally distinct phases of synaptic inhibition, namely, basal, early evoked, and sustained feedback synaptic inhibition in both GCs and SGCs and characterize the temporal profile and reliability of dentate evoked-sustained inhibition. We demonstrate that evoked-sustained feedback inhibition is not unique to GC but is present in SGCs as well. However, given the time dependent changes in evoked-sustained inhibition and robust summation of robust IPSC summation, we did not evaluate activity dependent changes in extrasynaptic GABA currents which are best examined in steady state. Although SGCs receive more frequent basal sIPSCs, GCs show greater enhancement of sustained feedback IPSCs upon afferent activation. These data identify that compared to GCs, SGCs exhibit blunted elevation in inhibitory gating following input activation, which likely facilitates their ability to sustain prolonged firing in response to afferent activation.

Although GCs and SGCs differ in their development, structure and excitability (Williams et al., 2007; Gupta et al., 2012; Save et al., 2018; Gupta et al., 2020), neurochemical markers to distinguish the cell types are lacking and morphology, characterized in rats, is used as to classify SGCs and GCs. Whether the neuropeptide proenkephalin (PENK), expressed in some prox-1 positive dentate neurons tagged based on increases in the immediate early gene FOS (Erwin et al., 2020), is selective for SGCs remines to be determined. However, our clustering analysis based on morphometric parameters, and the PCA of underlying features validate the use of number of primary dendrites (>3), dendritic angle, and soma-aspect ratio to distinguish SGCs from GCs. Moreover, dendritic complexity, which could influence intrinsic physiology and input integration (Gulledge et al., 2005; van Elburg and van Ooyen, 2010; van der Velden et al., 2012), was significantly lower in SGCs than in GCs, as reported in rats (Gupta et al., 2020). Curiously, putative GCs segregated into two subclusters, based on dendritic complexity, dendritic angle, and number of primary dendrites raising the possibility that physiology of dentate projection neuron subtypes scales with number of primary dendrites and dendritic complexity. Nevertheless, the physiological differences between GCs and SGCs identified here and in earlier studies (Williams et al., 2007; Larimer and Strowbridge, 2010; Gupta et al., 2012; Save et al., 2018) justify examination of their role in the dentate circuit.

Consistent with prior findings in rats (Gupta et al., 2012; Gupta et al., 2020), SGCs in mice receive more frequent sIPSCs than GCs. Since SGC somata and dendrites are located in the inner molecular layer, outside the dense axonal arbor of PV-INs in the granule cell layer, we anticipated fewer PV-INs inputs to SGCs than to GCs. Surprisingly, halorhodopsin suppression of PV-INs failed to reduce sIPSC frequency in both GCs and SGCs in spite of reliably hyperpolarizing (>5 mV) and reducing firing in YFP positive PV-INs. Moreover, channelrhodopsin activation of PV-INs evoked robust increases in GC and SGC IPSCs demonstrating viable, synaptically connected PV-INs. Furthermore, the same amber light protocol reduced evoked sustained IPSCs in SGCs, confirming the ability of halorhodopsin to suppress of PV-IN firing and synaptic release (Mahn et al., 2016). Why, then, is PV-IN suppression unable to reduce action potential dependent sIPSCs in GCs and SGCs even in the absence of glutamatergic blockers? One possibility is that PV-IN suppression disinhibits and increases firing of a subset of non-PV interneurons which compensate for the loss of PV-IN mediated inhibition. In this regard, the reciprocal activity states of PV and CCK neurons, recently reported in CA1, suggests that such a compensation is possible (Dudok et al., 2021). However, whether such a mechanism would exactly compensate for the reduction in PV-IN mediated IPSCs in a slice is debatable and cannot be resolved by using chemogenetic strategies which would potentially suffer from similar network level changes. The inability of optical/chemogenetic silencing to disambiguate direct from circuit-effects need to be considered while using cell-specific silencing strategies (Bernard, 2020). Interestingly, a prior study found limited contribution of PV-INs to action potential independent miniature IPSCs in granule cells (Goswami et al., 2012) and mutations impairing PV expression were shown to alter granule cell evoked IPSCs without impacting sIPSC frequency (Lucas et al., 2010). These findings are consistent with our results and raise the possibility that diverse populations of hilar and molecular layer interneurons may contribute substantively more than PV-INs to basal sIPSCs in GCs and SGCs. While our findings do not imply lack of PV-INs contribution to basal inhibition *in vivo*, the most parsimonious conclusion from our data is that PV-INs have limited role in basal sIPSCs in GCs and SGCs in slices.

PV-INs mediate robust and precise feedforward and feedback inhibition of GCs (Andersen et al., 1966; Hefft and Jonas, 2005; Ewell and Jones, 2010). We find that optical activation of PV-INs robustly increases IPSC frequency in both SGCs and GCs demonstrating that, despite their location in the inner molecular layer, SGCs receive inhibitory synapses from PV-INs. Moreover, afferent evoked IPSC peak amplitude and kinetics of were not different between GCs and SGCs indicating that, unlike spontaneous IPSCs, inhibition evoked by a single perforant path stimulus is comparable between the cell types.

Finally, we examined sustained feedback inhibition, previously found to correlate with SGC firing and advanced as a mechanism for lateral inhibition (Larimer and Strowbridge, 2010). We confirm that paired suprathreshold PP stimulation elicits persistent SGC firing in about 50% of trials and was accompanied by a two-fold increase in IPSC frequency in both GCs and SGCs. Sustained SGC activity followed the early evoke firing by over 200ms and persisted for over 10s. Interestingly, although individual SGC firing was observed only in half the trials, GCs and/or SGCs showed evoked sustained IPSCs in all slices examined, suggesting that activation of a subset of SGCs may be sufficient to sustain feedback inhibition. We report that about half the PV-INs examined showed increased firing following paired PP stimulation with latency and duration corresponding to that of evoked sustained inhibition in GCs and SGCs. Notably, even PV-INs lacking sustained firing showed increased EPSP frequency following paired PP stimulation indicating enhanced glutamatergic drive. Since SGC axons selectively innervate PV-IN somata (Rovira-Esteban et al., 2020), our findings are consistent with sustained firing in SGCs, which does not occur in GCs, driving sustained PV-IN activity.

Although both SGCs and GCs show sustained increase in IPSCs following paired afferent stimulation, the relative increase in IPSC frequency in SGCs is significantly lower than in GCs. Thus, SGCs which receive more robust polysynaptic NMDA currents than GCs (Larimer and Strowbridge, 2010) are, yet, subject to less gating feedback synaptic inhibition placing them in a privileged position to support persistent firing. This enhanced firing likely promotes expression of activity dependent genes and recruitment of SGCs as part of cellular ensembles during behavioral tasks (Erwin et al., 2020). Our analyses identify selective contribution of PV-INs to sustained feedback inhibition in SGCs but not in GCs. These results are paradoxical since wide field optical activation of PV-INs increases IPSCs in both SGCs and GCs and raises an intriguing possibility of target specific differences in PV-IN synaptic release and/or short-term plasticity similar to those reported in glutamatergic terminals (Sylwestrak and Ghosh, 2012; Blackman et al., 2013; Larsen and Sjostrom, 2015; Eltes et al., 2017) or selective reciprocal connectivity between SGCs and a subset of PV-INs.

In conclusion, our study demonstrates an input strength dependent shift in three distinct temporal phases of synaptic inhibition between GCs and SGCs. During steady-state basal activity, SGCs receive greater synaptic inhibitory drive than GCs. In response to afferent activation, evoked synaptic inhibition in SGCs appears comparable to GCs, which, given the greater glutamatergic inputs reported in SGCs (Larimer and Strowbridge, 2010), represents a relative reduction in scaling of inhibition to excitation in SGCs. With further increase in drive during paired stimulation, the increase in sustained feedback inhibition in SGCs is lower than in GCs. This activity dependent blunting of synaptic inhibitory inputs in SGCs compared to GCs indicates that SGCs, by having reduced activity dependent inhibitory gating, may support a distinct role in dentate processing by integrating, rather than sparsifying, inputs and driving lateral inhibition and selectivity of GCs.

## Acknowledgements

We thank Dr. Luke Fritzky at the Rutgers Imaging Core for help with imaging, Dr. Archana Proddutur for immunostaining validation of transgenic mouse lines and Dr. Deepak Subramanian for developing schematics. The project was supported by NIH/NINDS R01 NS069861, R01NS097750 and NJCBIR CBIR16IRG017 to V.S., NJCBIR CBIR15FEL011 to M.A and CBIR11FEL003 to A.G.

## Author Contributions

M.A., B.S performed experiments; M.A., A.G. and H.X. analyzed data; M.A., H.X., and V.S. interpreted results of experiments; M.A., prepared figures; M.A. and V.S. conceived of and designed research; M.A., and V.S. drafted manuscript.

